# Multi-omic profiling defines subtype-specific extracellular vesicle signatures in early human pregnancy

**DOI:** 10.64898/2026.03.10.710888

**Authors:** Kobe A. Abney, Taylor Hollingsworth, Alexandra Schneider, Elizabeth M. Brown, Hossein Fazelinia, Lynn A. Spruce, Rita Leite, Samuel Parry, Nadav Schwartz, Colin C. Conine, Rebecca A. Simmons

## Abstract

Early human pregnancy (11-15 weeks) is a critical period when placental dysfunction begins but remains clinically undetectable, preventing early intervention for pregnancy complications. This highlights a pressing need for circulating biomarkers that can signal the onset of a complication. Circulating extracellular vesicles (EVs) carry bioactive cargo that reflect originating cell activity in real-time, positioning them as early biomarkers of pregnancy complications. While pregnancy studies have identified plasma EV cargo alterations across complications, studying small EVs alone—the field standard—or bulk EVs together obscures pathological changes that may manifest preferentially in one population. To address this, we performed the first multi-omic comparison of small and large plasma EVs from healthy pregnancies at 11-13 weeks (n=10) to establish subtype-specific cargo differences and provide a reference for normal pregnancy EV biology. We quantified mitochondrial DNA (mtDNA), performed transcriptomics (mRNAs, miRNAs) and proteomics to define small and large EV signatures. Large EVs were preferentially enriched for mitochondrial cargo including DNA, transcripts, and proteins as well as hypoxia response, endocrine signaling, and VEGF signaling related transcripts and proteins. In contrast, small EVs were enriched for canonical EV markers and basement membrane proteins. Additionally, we found that EV miRNA composition was similar across EV subtypes with only 7 miRNAs differentially abundant between small and large EVs. These distinct small and large EV cargo signatures underscore the importance of profiling both populations for liquid biopsy potential. Together these data provide a baseline for future studies linking EV signature alterations to placental dysfunction and adverse pregnancy outcomes.

**Summary:** Size-fractionation and multi-omics reveal large plasma EVs in early pregnancy carry distinct mitochondrial, oxygen-sensing, and placental cargo missed by small EV-only or bulk EV biomarker studies.

## Introduction

The early pregnancy window between the first and second trimester (11-15 weeks) is a critical period for establishing healthy placental function, maternal vascular remodeling, and metabolic adaptation as the placenta transitions from a low oxygen to a relatively higher oxygen environment. When alterations in these processes arise during this transition the risk for placental dysfunction and subsequent adverse outcomes such as preeclampsia, preterm birth, and fetal growth restriction becomes increasingly high. A critical obstetric challenge is that the etiology of these complications are largely unknown and clinical signs often only manifest late in pregnancy, limiting intervention and personalized care. Identifying molecular signatures of placental dysfunction during this early window could enable better risk assessment and targeted intervention before irreversible damage occurs.

Circulating extracellular vesicles (EVs) represent a promising source of minimally invasive biomarkers for monitoring placental health and proper pregnancy progression in real-time. Throughout gestation, both maternal and placental cells release EVs that carry functional nucleic acids, proteins, and metabolites into maternal circulation as a form of maternal-fetal communication [1]. EVs not only reflect the cellular activity of the originating cell through the types of cargo enclosed but also carry bioactive cargo that alters recipient cellular functions. For example, EV RNAs have been shown to regulate gene expression and translate functional proteins, while EV proteins modulate intercellular signaling [2–8]. Moreover, EV mitochondrial cargo is widely profiled across disease states to assess mitochondrial dysfunction and systemic metabolic activity. Specifically, intact mitochondria and mtDNA in EVs have been shown to incorporate into the recipient cells’ mitochondrial network and rescue metabolically impaired cells, respectively [9–13]. Accumulating evidence suggests EVs promote successful pregnancy by mediating key physiological processes such as placental metabolism, implantation, immunomodulation, spiral artery remodeling, and labor [14–16]. Importantly, changes in EV concentration and cargo have been described across many pregnancy complications, highlighting the capacity of EVs to reflect maternal and fetal tissue dysfunction and potentially indicate the risk of pregnancy complications [15,17].

Despite the growing interest in EV-based biomarkers, most EV pregnancy studies have focused predominantly on small EVs (sEVs; ∼50–150 nm) or examined bulk EV populations leaving large EVs (lEVs; ∼150–500 nm) relatively underexplored despite their potential for distinct molecular cargo and biological insights [15]. These two subtypes arise from distinct biogenesis pathways, positioning each subtype to carry fundamentally different molecular information that may reflect complementary aspects of placental and maternal adaptation during early pregnancy. In cell culture systems, sEVs originate predominantly from inward budding of multivesicular bodies via endosomal sorting complexes (ESCRT) and lipid or tetraspanin-mediated mechanisms, enriching for small RNAs and proteins reflective of intracellular signaling pathways [18–25]. In contrast, lEVs arise from outward budding of the plasma membrane through processes regulated by lipid dynamics and cytoskeletal remodeling, enriching for cargo that reflects the plasma membrane and cellular responses to stress or signaling cues [26–31]). Although distinct biogenesis mechanisms exist in controlled systems, circulating EVs in complex biofluids like plasma likely represent a heterogenous population derived from many different cell types employing multiple biogenesis pathways and cargo sorting mechanisms [32,33]. This is a critical issue for identifying early biomarkers of placental dysfunction, as pathological changes may manifest preferentially in one EV subtype but not another. Therefore, size-based fractionation is essential to separately profile lEV and sEV populations and determine whether these subtypes carry distinct molecular information that, together, provide a more complete picture of early pregnancy health than either population alone.

To date, no study has comprehensively compared plasma-derived sEVs and lEVs during early gestation in healthy pregnancies across multiple molecular dimensions. This represents a critical knowledge gap, as single-omic approaches fail to reveal which EV subtype carries the most comprehensive cargo for detecting placental dysfunction or inadequate maternal adaptations. In this study, we perform size-fractionated molecular profiling of plasma EVs from normal pregnancies (11-15 weeks), using a multi-omic approach focusing on four biologically informative analytes: mitochondrial DNA (mtDNA), large RNAs (messengerRNA/mRNA and long non-coding RNA/lncRNA), small RNAs specifically microRNA/miRNA, and proteins. These analytes were selected to capture complementary signatures: mtDNA and mitochondrial proteins report metabolic state during the critical oxygen transition, large RNA transcripts reveal cellular activity and tissue origins, proteins modulate intercellular signaling activity in distant tissue, and regulatory miRNAs demonstrate capacity to alter recipient cell gene expression. Importantly, convergence of these layers would reveal enhanced pathway signaling from the originating cell that could serve as a baseline for future biomarker development. By establishing a reference profile for plasma EV subtypes from normal pregnancy, this work provides fundamental insight into how different EV subtypes contribute to maternal-fetal communication and metabolic adaptation, enabling future studies to identify pathological alterations in pregnancy complications for biomarker discovery.

## Methods

### Participant Cohort

The Placenta 3D study, a prospective longitudinal cohort, is an ongoing Institutional Review Board (IRB) approved study (IRB#852638) at the University of Pennsylvania. Participants are recruited from the Obstetrics & Gynecology (ObGyn) practice at the Hospital of the University of Pennsylvania (HUP). Participants presenting for a prenatal ultrasound prior to 15 weeks and 6 days’ gestation at the Maternal Fetal Medicine Ultrasound unit at HUP were screened for eligibility through medical record review. Eligible patients are approached for study participation by a Clinical Research Coordinator. Inclusion criteria include a singleton pregnancy, maternal age of 18 years or older, ability to read and understand English, and willingness and ability to provide written Informed Consent. Exclusion criteria include multiple gestation and the presence of a known major fetal anomaly or genetic abnormality.

### Clinical and Demographic Data Collection

Clinical and demographic information including maternal age, gestational age at sample collection and delivery, maternal body mass index (BMI), birthweight, fetal sex and race was abstracted from the electronic medical record (Table 1). Samples were collected during Visit 1 appointments occurring within a 11-15 week window per study protocol. For this study, we sampled ten normal pregnancies, with gestational ages ranging from 11.1 to 13.4 weeks, uncomplicated by adverse outcomes such as preterm birth, preeclampsia, or fetal growth restriction. BMI of the women in the study ranged between 24-30 kg/m^2^, representative of the average BMI of the total cohort.

**Table 1.** Cohort Demographics. Samples were collected at Visit 1 appointments (11-15 wks). Maternal age, gestational age, birthweight, and BMI are represented as mean ±(SD). Gestational age and birthweight are followed by range.

### Platelet-poor Plasma Preparation for EV Isolation

Blood was collected from a maternal peripheral vein at the first visit (V1) appointment in a K2EDTA tube. Blood was inverted gently and spun at 1000*g* for 10 minutes at room temperature to isolate plasma. 1 ml aliquots of plasma were transferred to 1.7 ml microcentrifuge tubes and spun at 2500*g* for 15 minutes at room temperature to deplete cellular debris and platelets. Aliquots were transferred to new 1.7 ml tubes and spun again at 2500*g* for 15 minutes at room temperature [34–38]. The platelet-poor plasma (PPP) aliquots were transferred to cryovials (Fisherbrand, Waltham, Massachusetts, Cat # 10-500-26) and stored at -80°C.

### Isolation of Large EVs by Centrifugation

Centrifugation was performed to enrich lEVs (200-500 nm) from human PPP. For each subject, samples for mRNA analysis were derived from a 1 ml aliquot. 800 µl of a second aliquot was used for DNA isolation, and 1ml of a third aliquot was used for proteomic analysis. PPP samples were taken from -80°C storage and thawed overnight at 4°C for experiments. Using an Eppendorf 5424 tabletop centrifuge (Eppendorf, Enfield, Connecticut), plasma samples were spun at 20,000*g* for 30 minutes at room temperature. The supernatant was removed and transferred to a new Eppendorf tube for size exclusion chromatography. The remaining pellet (crude lEVs) was washed with 1 ml of filtered 1X PBS and centrifuged at 20,000*g* for 30 minutes at room temperature to remove any remaining debris. The supernatant was discarded, and the crude lEV pellet was resuspended in 78.4 µl of filtered 1X PBS for RNA extraction or 100 µl for mtDNA extraction. 15 µl of sample was aliquoted for particle counting.

### Isolation of Small EVs by Size Exclusion Chromatography and Ultracentrifugation

Size exclusion chromatography was performed to enrich sEVs using the qEV1 Gen 2 70 nm column (Izon Science, Christchurch, New Zealand) and the Automatic Fraction Collector (Izon Science, Christchurch, New Zealand) as described for plasma EVs. The supernatant collected from the first 20,000*g* spin was used on the column. Of the four eluted fractions, fractions 1-3 were combined and concentrated at 100,000*g* for 70 minutes using the Optima Max-TL Ultracentrifuge and TLA 110 rotor (Beckman Coulter, Indianapolis, Indiana) at 4°C to yield a concentrated sEV pellet. The supernatant was discarded and sEVs were resuspended in 78.4 µl of filtered 1X PBS for RNA extraction or 100 µl for mtDNA extraction. 15 µl were aliquoted for particle counting.

### Nanoparticle Tracking Analysis and Cryo-electron Microscopy

EV were characterized by Nanoparticle Tracking Analysis (NTA), cryo-electron microscopy (cryo-EM), and protein surface markers per MISEV23 guidelines (Welsh, 2023). All NTA measurements were done in triplicates using the ZetaView (Particle Metrix, Inning am Ammersee, Germany) at 25°C, running on ZetaView analysis software v8.05.14 Sp7. Each EV sample was diluted in DI water directly before measurement based on manufacturer’s recommendations of 140-200 particles/frame. lEV measurements were acquired using the following parameters: 520 nm laser, scatter filter wavelength, 25 min to 255 max brightness, tracking radius 100, camera-[Fp Sec 30, 1 cycle, frame rate 7.5], shutter 100, and sensitivity 70. sEV measurements were acquired using: 520 nm laser, scatter filter wavelength, 20 min to 255 max brightness, tracking radius 30, camera-[Fp Sec 60, 1 cycle, frame rate 30], shutter 100, and sensitivity 80. All measurements were captured in eleven fields.

For cryo-EM, 5 µl of plasma EV sample was applied onto the carbon side of the EM grid which was then blotted for 2.0 seconds and plunge-frozen into liquid ethane and imaged with a 300 keV Titan Krios G3i microscope, featuring a phase plate, K3 Summit direct detector camera, and BioQuantum GIF energy filter.

### Immunoblotting

Three lEV and three sEV pellets were resuspended in 30 µl of electrophoresis buffer (40 mM Tris-HCl pH 6.8, SDS 5%, 2M Thiourea, 6M Urea, 0.1 mM EDTA, blue bromo phenol traces) and incubated overnight at room temperature. CD9 expression was measured under non-reducing conditions, while apolipoprotein B-100 (apoB-100) expression was assessed under reducing conditions. The total volume (30 µl) of each sample was loaded per lane and run on Mini-Protean TGX precast 4-20% gels (Bio-Rad, Hercules, California) with Precision Plus Dual Color ladders ((Bio-Rad, Hercules, California) at 120V for 1 h. Proteins were transferred onto premium nitrocellulose membranes (Cytiva, Freiburg im Breisgau, Germany) using 200 mA over 3 h on ice. After blocking with 100% Intercept Blocking Buffer (LiCor, Lincoln, Nebraska) at room temperature for 2 h, the membranes were washed in 0.15% TBST and incubated overnight at room temperature with primary antibodies in 50% Intercept Blocking Buffer. The following primary antibodies were used: anti-CD9 (biotinylated clone HI9a, 1:1000 w/v dilution, Biolegend Cat # 312112) and anti-APOB (clone C1.4, 1:1000 w/v dilution, Santa Cruz Cat # 13538). The membranes were washed in 0.15% TBST and incubated with secondary antibodies pre-conjugated with IR-dye: Streptavidin (LiCor Bio Cat # 926-32230) and goat anti-mouse (LiCor Bio Cat # 926-32210) diluted 1:5000 in 0.15% TBST at room temperature for 2 h. The blots were imaged with the LiCor Odyssey Imager (LiCor, Lincoln, Nebraska).

### Ethanol precipitation-based DNA extraction

Some EV-mtDNA studies have included a DNase treatment step during the DNA isolation process to degrade unencapsulated DNA. However, this step disturbs the EV corona, a dynamic layer of proteins and nucleic acids adhering to the surface of EVs, which has been shown to play an essential role in EV functionality, influencing angiogenesis, cell uptake, and immunomodulation [39]. Moreover, removal or disturbance of the corona, through additional purification or enzymatic treatments can significantly alter EV functions, including their uptake by recipient cells and their biological activity [40]. Therefore, here, EVs were not DNase-treated to preserve the corona and capture biologically active, EV-associated mtDNA content in its native, unstripped state.

We adapted a previously reported method used to isolate cell-free mtDNA [41]. EV-associated mtDNA was extracted from volumes equal to 2 x 10^7^ lEVs or 1 x 10^8^ sEVs and diluted with PBS to a final volume of 37 µl. EVs were incubated at 55°C for 16 h with 1.66 µl of 10% SDS, 37.9 µl of 1 M Tris-HCl (pH 8.5), 7.6 µl of 0.5 M EDTA, 3.12 µl 5 M NaCl, 2.62 µl of 20 mg/ml Proteinase K, and 0.70 µl of 1% β-mercaptoethanol. Samples were digested for 16 h, cooled at room temperature for 5 minutes, and then diluted 4.5:1 with a premixed digestion buffer containing 0.2% SDS, 100 mm Tris-HCl, 5 mm EDTA, 200 mm NaCl based on the concentration of the stock solutions described above. Following digestion, samples were incubated for 5 minutes on ice with 170 µl of 5 M NaCl to precipitate proteins. The samples were then centrifuged at 21,000*g* for 20 minutes at 4°C. The supernatant was transferred to a new tube and 800 µl of cold 100% ethanol and 1 µl of GlycoBlue were added. The tubes were centrifuged again at 21,000*g* for 20 minutes at 4°C. The supernatant was aspirated, and the remaining pellet was washed with 500 µl of cold 70% ethanol and centrifuged at 21,000*g* for 20 min at 4°C. The supernatant was aspirated again, and the pellet air-dried at 37°C for 15 minutes. 60 µl of TE buffer containing 2% RNase A (200 µg/ml) was added to the tubes and incubated at 55 °C for 1 h. The resuspended DNA was aliquoted and stored at −20 °C.

37 µl of EV-depleted samples were used to isolate mtDNA as a control following the protocol above. EV-depleted samples were derived from the remaining supernatant after both lEVs and sEVs were pelleted.

### EV-mtDNA Measurement

EV-associated and EV-depleted mtDNA was quantified using a TaqMan-based quantitative polymerase chain reaction (qPCR). These reactions were run in triplicate on a QuantStudio 7 real-time PCR system (Thermo Fisher Scientific, Waltham, Massachusetts). As previously described, the assay targeted mitochondria-encoded human NADH:ubiquinone oxidoreductase core subunit 1 (*MT-ND1*) [41,42]. Thermocycling conditions were as follows: an initial denaturation step at 95 °C for 20 s followed by 40 cycles of 95 °C for 1 s, 63 °C for 20 s, and 60 °C for 20 s.

While the CT value can indicate the relative presence of mtDNA, it does not quantify absolute copy number present in the sample. Therefore, serial dilutions of pooled human placental DNA quantified for *MT-ND1* (copies per microliter) by digital droplet PCR were used to create a standard curve [42]. Using the standard curve, we first calculated the abundance of mtDNA present. We then normalized the abundance by the number of EVs in the starting material to estimate the number of mtDNA copies per EV [42]. While estimating mtDNA copy number per EV is an oversimplification given the inherent heterogeneity of EV populations, this approach provides a necessary framework for direct quantitative comparison of mtDNA content between sEVs and lEVs.

### Total RNA Extraction

As previously described [43], sEV and lEV samples (resuspended in 63.4 uL of PBS) were diluted to 100 µL by the addition of 33.3 µL lysis buffer (6.4 M Guanidine HCl, 5% Tween 20, 5% Triton, 120 mM EDTA, and 120 mM Tris pH 8.0) and 3.3 µL Proteinase K, then incubated at 60°C for 15 minutes while shaking to extract total RNA. 100 µL of water and 200 µL of TRI reagent (Invitrogen, Waltham, Massachusetts, Cat # AM9738) were added to the sample and mixed thoroughly to ensure complete lysis. The samples were transferred to a phase lock tube (discontinued; Quantabio, Beverly, Massachusetts) that was previously centrifuged empty at 20,000*g* for 1 minute. BCP (1-bromo-2 chloropropane) was added to the phase lock tube at one-fifth of the total volume (80 µL) and inverted 15 times to mix. The samples were centrifuged at 4°C for 4 minutes at 20,000*g*, and the aqueous phase was transferred to a new low-adhesion tube. The aqueous phase volume was measured for each sample before adding 1 µl of Glycoblue (Invitrogen, Waltham, Massachusetts, Cat # AM9516) and 1.1 times the volume of isopropanol. Samples were mixed thoroughly and incubated for 1 h at -20°C to precipitate the RNA. Following precipitation, RNA was pelleted at 20,000*g* for 15 minutes at 4°C and washed with freshly made, ice-cold 70% ethanol. Following the wash, the RNA pellet was reconstituted in 87.5 µL RNase-free water and incubated with 10 µL RDD buffer and 2.5 µL DNase I (Qiagen, Hilden, Germany, Cat # 79254) for 15 minutes at room temperature. 100 µL of water and 400 µL of TRI reagent was added to the sample and mixed thoroughly. The samples were transferred to phase lock tubes for phase separation and isopropanol precipitation, as described above. The final RNA pellets were reconstituted in RNase-free water for low-input mRNA library preparation.

### Low-input RNA Library Prep and Next Generation Sequencing

Long RNA Libraries were prepared following an adapted protocol from single-cell RNA seq using the SMART-Seq protocol [43,44]. Polyadenylated RNAs from sEVs and lEVs were reverse transcribed into cDNA using Superscript II (Invitrogen, Waltham, Massachusetts, Cat# 18064014). The cDNA was amplified with 20 PCR cycles and cleaned up with AMPure XP beads (Beckman Coulter, Indianapolis, Indiana, Cat# A63881). cDNA was then quantified with the Qubit dsDNA HS Assay Kit (Life Technologies, Waltham, Massachusetts, Cat# Q32851). Using the Nextera XT kit (Illumina, San Diego, California, Cat# FC-131-1096), a pool of uniquely indexed samples was constructed from 2 ng of the purified PCR product. A final 12 cycle amplification was performed on the pooled libraries and cleaned up using the AMPure XP beads for sequencing on a NextSeq1000.

Small RNA libraries were prepared using a modified SMARTer® smRNA-Seq Kit for Illumina (Takara Bio USA, San Jose, CA) optimized for low-input small RNA species including miRNA, small interfering RNA (siRNA), PIWI-interacting RNA (piRNA), transfer RNA (tRNA), and small nucleolar RNA (snoRNA). RNA from EVs was polyadenylated and reverse transcribed into cDNA using PrimeScript RT (Takara Bio USA, San Jose, CA) with template switching to incorporate adapter sequences. To account for differences in starting material, representative samples were used to optimize PCR amplification and determine the minimum number of cycles required without overamplification. Based on this optimization, libraries from sEV RNA were amplified with 20 PCR cycles and libraries from lEV RNA were amplified with 18 PCR cycles using uniquely indexed primers. Amplified libraries were purified using AMPure XP beads (Beckman Coulter, Indianapolis, Indiana) and resolved on an 8% polyacrylamide gel. The small RNA library fraction corresponding to 171–193 bp was excised to remove adapter dimers and nonspecific products. Purified libraries were quantified with the Qubit dsDNA HS Assay Kit (Life Technologies, Thermo Fisher Scientific, Carlsbad, CA). Using the Nextera XT kit (Illumina, San Diego, CA), a pool of uniquely indexed samples was constructed from 2 ng of each purified library. The pooled libraries were prepared with a 5% PhiX spike-in and sequenced single-end on a NextSeq 1000. Adaptor sequences and 5’ adenylation from library preparation were removed using Cutadapt.

### Protein Extraction and Digestion

sEV and lEV pellets were lysed in 8 M urea, 5% SDS, 100 mM Tris-HCl (pH 8), and protease inhibitors, followed by benzonase treatment. Protein concentration was estimated by in-gel Bradford staining using an E. coli lysate standard.

Proteins were digested using the S-Trap Micro protocol. Samples were reduced (TCEP), alkylated (iodoacetamide), acidified, and digested with trypsin/Lys-C. Peptides were eluted, vacuum-dried, desalted using C18 StageTips, and reconstituted in 0.1% TFA containing iRT peptides. Peptide concentration was measured at OD280 and adjusted to 400 ng/µL for injection.

### Mass Spectrometry Acquisition

Samples were randomized and analyzed on a Bruker TimsTOF HT coupled to a NanoElute2 nano-LC in DIA-PASEF mode. Peptides (800 ng) were separated on a 25 cm C18 column using a 35-minute gradient (2–30% acetonitrile). Data were acquired over an m/z range of 100–1700 with ion mobility separation (1/K0 0.70–1.30 V·s/cm²). DIA windows covered 299.5–1200.5 Da with a cycle time of 0.65 s. Ion mobility was calibrated using Agilent tuning mix standards.

### Proteomic Quality Control and Data Processing

Instrument performance was monitored using QuiC software with spiked-in iRT peptides. A K562 digest standard was injected before and after the sample set for system suitability assessment. Raw files were processed in Spectronaut (v20.2) using directDIA against the human UniProt reference proteome with common contaminants appended. Trypsin specificity (≤2 missed cleavages), carbamidomethylation (fixed), and N-terminal acetylation and methionine oxidation (variable) were specified. Peptide and protein FDR were controlled at 1%.

### Bioinformatic Analysis

Statistical analysis was performed using GraphPad Prism10. Outliers were determined using the ROUT method with the FDR(Q)=1% in Prism10. MtDNA samples with values below the limit of detection were excluded prior to statistical analysis. Normality of data was determined using the Shapiro Wilk test (α = 0.05). Following this, for normally distributed data an unpaired two-tailed Student’s t-test was performed to assess the difference of two means. Significance was defined as p < 0.05. Cohort demographics is presented as mean values ± standard deviation.

Long RNA data were mapped to the hg19 reference genome using Feature Counts on Dolphin Next and normalized to transcripts per million (TPM). Sequencing quality control was assessed by determining the number of transcripts within each sample with TPM ≥ 1 [45]. lEV samples with less than 2,100 and sEV samples with less than 550 genes satisfying this cutoff were removed. Three lEV samples and two sEV samples did not meet this cutoff and were excluded from downstream RNA-seq analyses. Raw read counts were uploaded to R Statistical Software and using the DESeq2 package data was normalized, and differential gene expression was determined using a false-discovery rate p-value [46]. To filter out low-abundant transcripts, only genes with at least 10 counts across four samples in either group were analyzed. Differentially abundant transcripts were determined using a log_2_FC >|1|, and an adjusted p-value <0.05. Ingenuity Pathway Analysis was used for functional enrichment data analysis of DEGs.

Small RNA bioinformatic pipelines were assembled using DolphinNext [45]. Read quality was assessed using FastQC, and reads were mapped sequentially to rRNA, miRbase, tRNAdb, snRNA, mRNA and Refseq using STAR and totaled using Feature counts. Mapped reads were normalized to total genome mapping reads for subsequent analysis. Small RNA differential expression was determined by a log_2_FC >|2| and an adjusted p-value <0.05 using DESeq2.

MS2 intensities were log2-transformed and median normalized in R. Proteins with complete data in at least one cohort were retained. We applied a two-level normalization strategy to ensure robust and comparable proteomics data. First, during sample preparation, all samples were processed together in a single batch and total protein concentrations were measured to normalize input amounts across samples, minimizing variability due to sample loading. Second, during data analysis, protein intensities were log2-transformed and median-centered so that each sample had a median intensity of zero. This approach reduces technical variability while preserving relative differences in protein abundance across samples. Differential abundance between groups was assessed using Limma, with significance defined as adjusted P value < 0.1 and log2FC > |1|. Proteins with statistically different abundance were prioritized for downstream bioinformatics analyses and visualized using volcano plots.

## Results

### Small and Large EV Characterization

sEV and lEVs were isolated from the plasma of pregnant women (n=10) and characterized following the Minimal information for studies of extracellular vesicles (MISEV) criteria (Figure 1a) [36]. Cryo-electron microscopy showed intact vesicles with the expected morphology of sEVs and lEVs (Figure 1b). The size distribution of two representative EV subtypes was confirmed by NTA, with sEVs peaking at approximately 100 nm and lEVs peaking at 200 nm (Figure 1c) (Supplemental Table S1 and Supplemental Table S2). Immunoblotting showed enrichment of CD9, an established EV marker, in the EV fractions compared to plasma (Figure 1d). While apolipoprotein B-100 (ApoB-100), a marker of lipoprotein contamination, was not completely depleted in the EV fractions, it was significantly reduced compared to intact plasma (Figure 1e). The presence of additional canonical EV markers such as CD63, CD81, and programmed cell death 6 interacting protein (PDCD6IP; ALIX) was confirmed through proteomic analysis. Together these data validate the presence of EVs in the samples that were used for downstream mtDNA, transcriptomic, and proteomic analysis.

**Figure 1.**
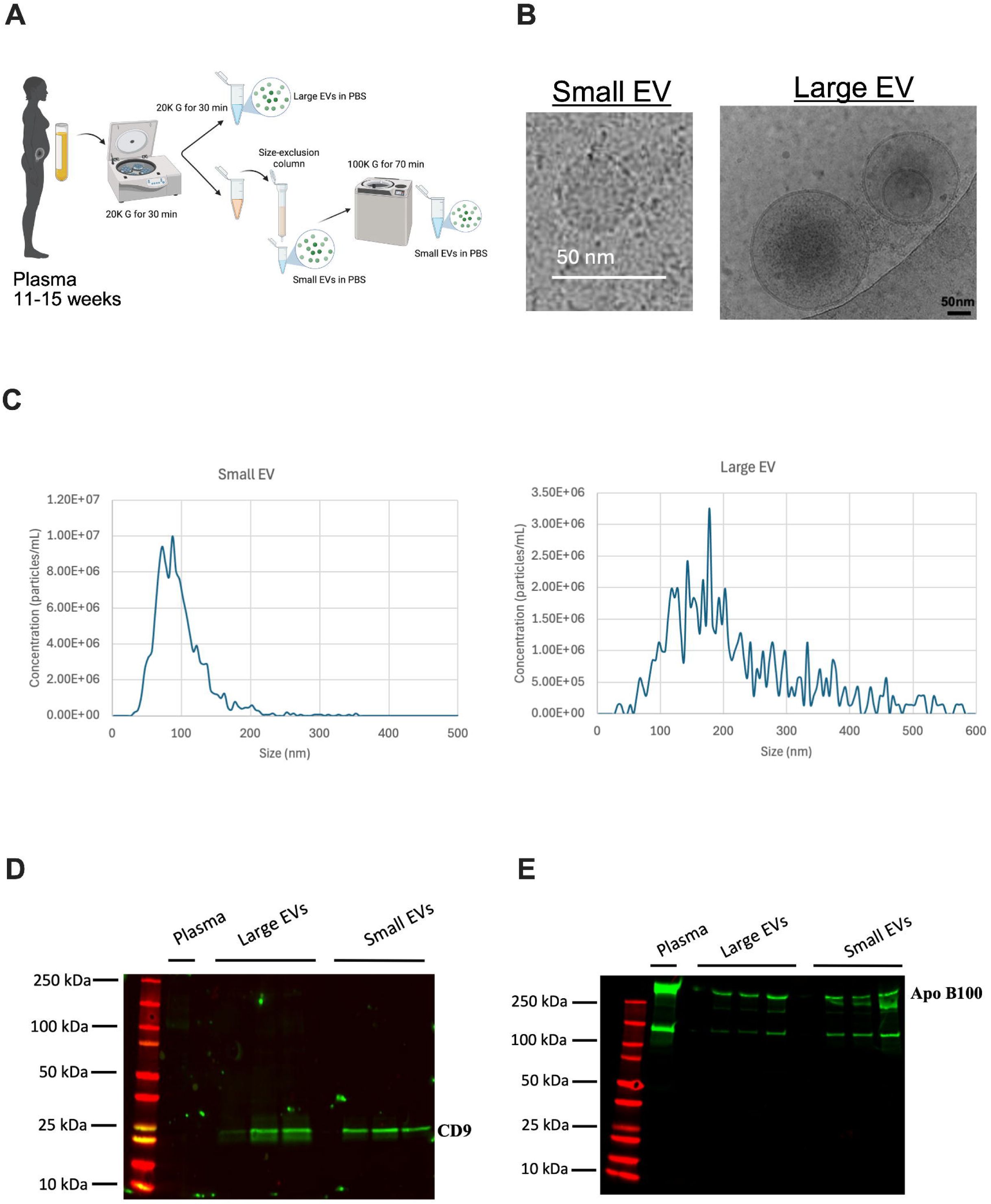
Extracellular vesicles were characterized by cryoEM, western blot, and nanoparticle tracking analysis. (A) EVs were isolated from first-trimester platelet-poor plasma via differential centrifugation and size exclusion chromatography. (B) Cryo-electron microscopy was used to visualize sEV and lEV morphology and size. (C) Nanoparticle tracking analysis was used to determine the size distribution of both sEVs and lEVs. Graphs represent one biological samples per EV subtype. (D) Plasma and plasma EV samples were immunoblotted with an antibody against CD9. (E) Plasma and plasma EV samples were immunoblotted with an antibody against lipoprotein Apo B100. Created in BioRender.

### Large EVs From Pregnant Women Are Associated with Higher Levels of mtDNA

mtDNA is a well-documented bioactive proxy for cellular energy metabolism and mitochondrial function. The quantity of mtDNA associated with EVs is indicative of cellular stress, highlighting its use in assessing systemic metabolism during early pregnancy [47]. To date, no studies have directly quantified and compared sEV- and lEV-associated mtDNA copy number during pregnancy. Therefore, we wanted to first compare the levels of EV-associated mtDNA in sEVs and lEVs from the plasma of pregnant women using qPCR. We observed that the mtDNA CT values from the lEV samples were significantly lower (p<0.001) than those of the sEVs (Supplemental Figure S1) indicating higher mtDNA content in lEVs. Quantifying absolute copy number, lEVs were associated with significantly more mtDNA copies compared to sEVs (Figure 2) (Supplemental Table S3).

**Figure 2.**
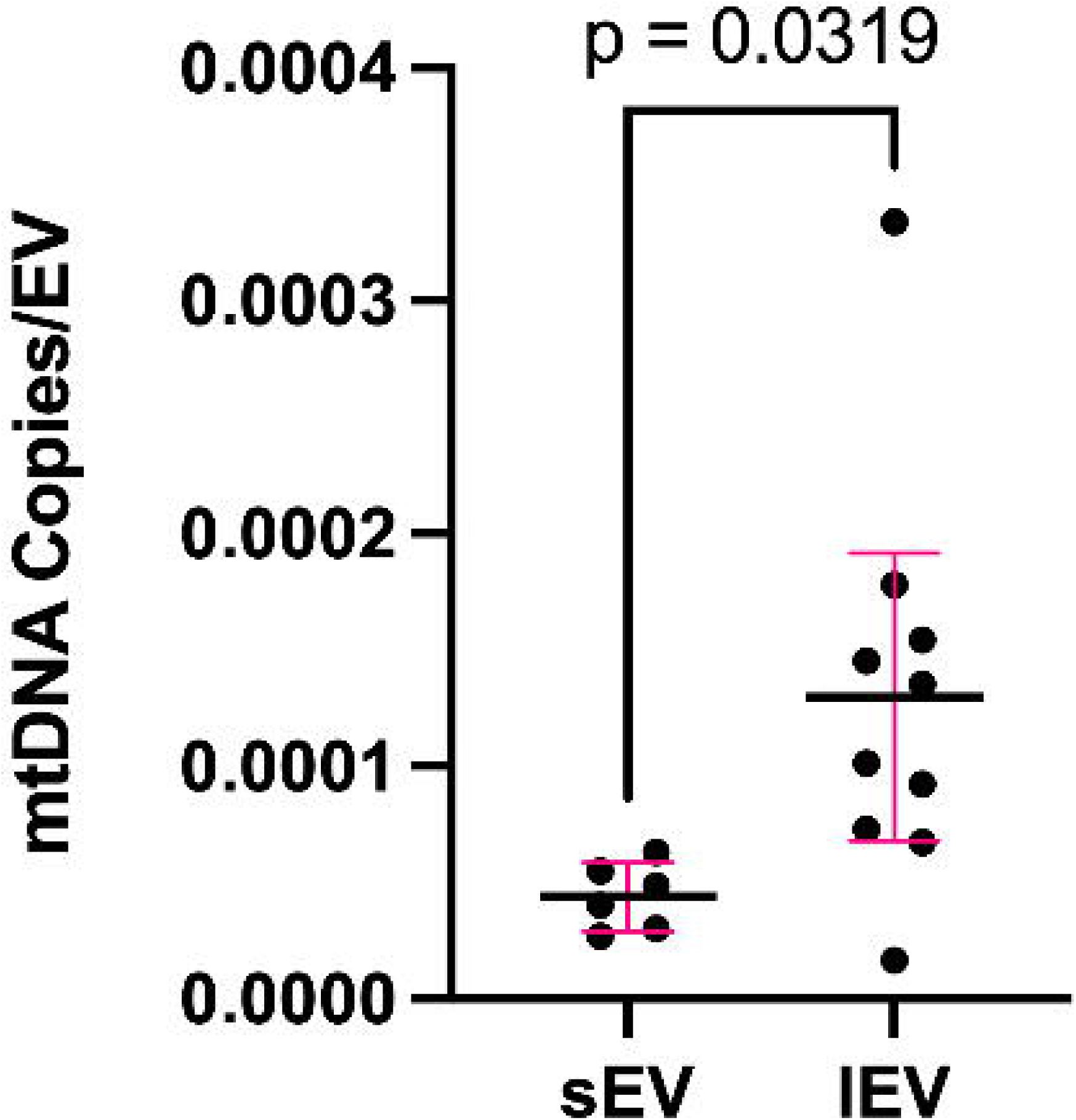
Early pregnancy large EVs are associated with more mtDNA. EV-associated mtDNA was quantified from sEVs and lEVs from plasma of pregnant women using qPCR. mtDNA copy number was calculated using a standard curve derived from human placenta mtDNA. An unpaired student’s t-test was performed to compare the number of mtDNA copies per sEV and lEV from 800 uL of platelet-poor plasma.

We quantified mtDNA in EV-depleted samples as a control for non-EV associated mtDNA. EV-depleted fraction showed near-absent mtDNA signal, indicating that the mtDNA detected in our preparations are EV-associated rather than freely circulating or attributable to intact mitochondria (Supplemental Figure S1).

### Long RNAs Are More Abundant in Large EVs from Pregnant Women

While sEV-miRNAs are well characterized across different disease states, characterization of EV-long RNAs during early pregnancy remains unexplored [2,21,24]. Here we isolated total RNA to quantify mRNA and lncRNA associated with sEVs and lEVs during early pregnancy.

To define subtype-specific RNA composition we first assessed the percentage of RNA reads mapping to mRNA and rRNA in each EV subtype. Similar to other non-pregnancy EV-RNA studies, we observed that lEVs were relatively enriched for mRNAs, whereas sEVs showed a higher proportion of rRNAs (Figure 3a) [3,48–52]. To our knowledge, this is the first description of such long RNA-class partitioning between size-defined plasma EV subtypes in pregnancy. Principal component analysis (PCA) separated samples by EV subtype, with lEVs clustering much tighter than sEVs (Figure 3b). A scatter plot using the average variance stabilizing transformation (VSD) counts, determined by the vst function in DESeq2, highlighted the disproportionate distribution of transcripts between lEVs and sEVs (Figure 3c) [46]. Overall, there were significantly more transcripts detected in the lEVs compared to sEVs.

**Figure 3.**
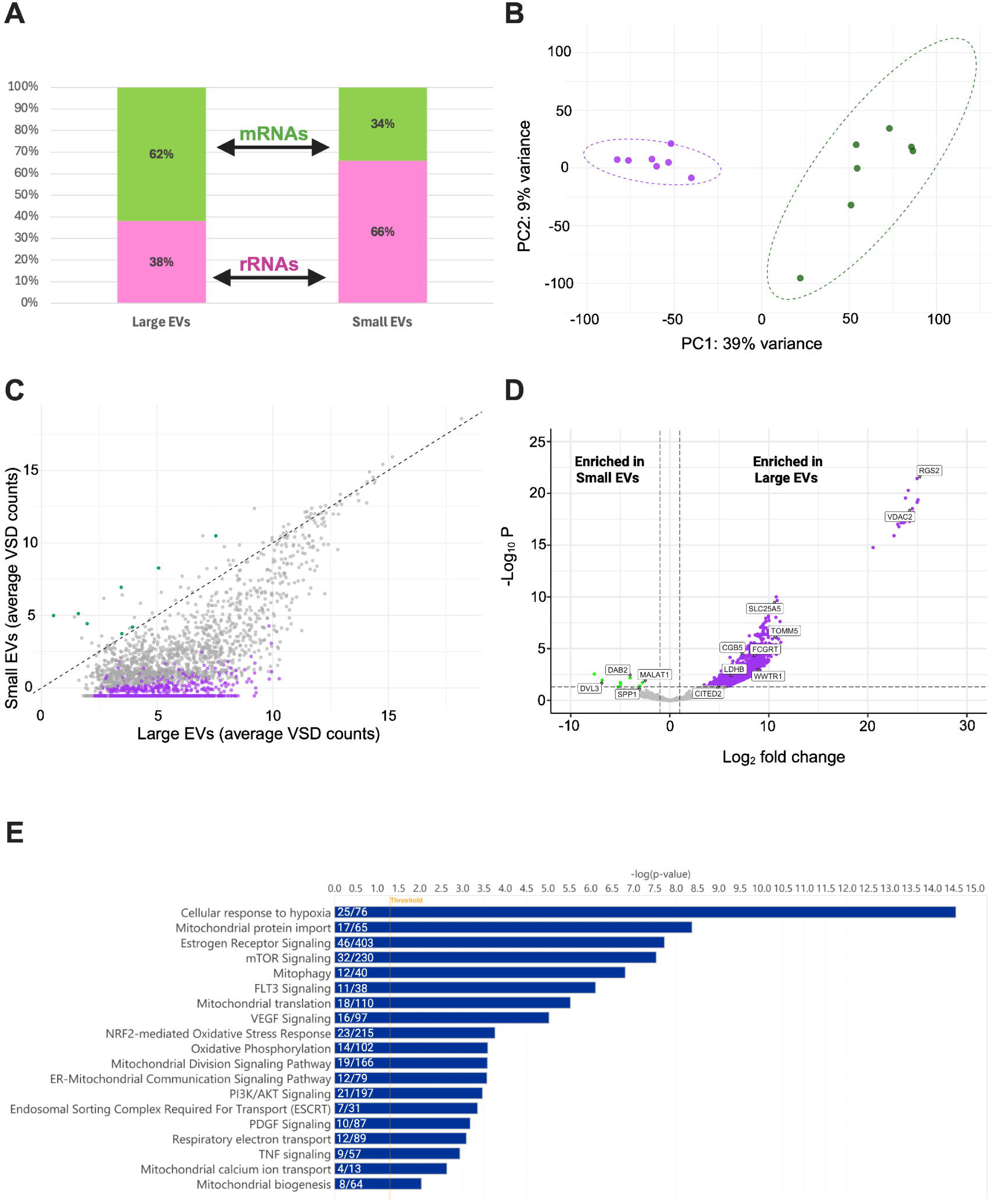
Early pregnancy large EVs are enriched with relevant long RNAs. RNA was extracted from matched sEVs and lEVs and long RNAs were sequenced using Next Generation Sequencing. (A) Percentage RNA class partitioning against total reads by EV subtype. Pink represents rRNAs and green represents mRNAs. (B) Principal component analysis (PCA) of the transcriptome of individual sEV and lEV samples isolated from maternal plasma (11-15 weeks gestation). In total, there were seven lEV samples and nine sEV samples used for downstream analysis. (C) Scatter plot of average variance stabilizing transformation (VSD) counts for lEVs (x-axis) versus sEVs (y-axis). Colored dots highlight long RNAs differentially abundant between EV subtypes. Purple indicates long RNAs enriched in lEVs and green indicates long RNAs enriched in sEVs. The grey transcripts highlight those that were not determined to be differentially abundant by DESeq2 in R. (D) Volcano plot depicting log_2_ fold change and -log_10_ adjusted *p*-value of identified transcripts in sEVs and lEVs. Colored dots indicate differentially abundant transcripts as determined using DESeq2. Significance threshold: log_2_FC ≥ |1| and adjusted *p*-value of ≤ 0.05. (E) Pathway enrichment of the differentially abundant transcripts between sEVs and lEVs using Ingenuity pathway Analysis. Ratio of genes in our dataset that overlap the total genes in the IPA database are represented for each pathway within the bar. Created in BioRender.

### Placental Transcripts Are Enriched in Plasma-Derived EVs

Placenta-derived extracellular vesicles and their cargo are of particular interest for understanding maternal-fetal communication in normal pregnancy physiology and pathophysiology of pregnancy complications. Prior studies have characterized placenta-derived EVs and distinguished cell-type origin using tissue markers to identify extravillous trophoblast, cytotrophoblast, and syncytiotrophoblast [14, 53–56]. In our dataset, we detected transcript level chorionic somatomammotropin hormone 1 (*CSH1*) and pregnancy specific beta-1-glycoprotein 1 (*PSG1*), both which are hallmarks of syncytiotrophoblast, as well as glycoprotein hormones alpha polypeptide (*CGA*) (Supplemental Table S4). Although the levels of these transcripts were not differentially abundant between sEVs and lEVs, their presence in both fractions are consistent with placental contribution to the circulating EV pool.

Using DESeq2, we then determined differentially abundant transcripts between sEVs and lEVs to establish EV subtype-specific signatures. To contextualize these signatures, we cross-referenced differentially abundant EV transcripts with published human placental transcriptomic data [57]. Out of the total 2688 genes with quantitated RNAs, 1142 were differentially abundant with the overwhelming majority enriched in lEVs (Figure 3d)(Supplemental Table S4). Only eight transcripts were more abundant in sEVs including one lncRNA, metastasis associated lung adenocarcinoma transcript 1 (*MALAT1),* and seven protein-coding transcripts. Of the protein-coding transcripts enriched in sEVs, secreted phosphoprotein 1 (*SPP1)*, DAB adaptor protein 2 (*DAB2)*, disheveled segment polarity protein 3 (*DVL3)*, trichohyalin (*TCHH*), and proline rich coiled-coil 2C (*PRRC2C*) have been identified in high abundance (FPKM >10) in human placental transcriptomic datasets [57]. Of the 1134 transcripts enriched in lEVs, 383 were also highly expressed (FPKM > 10) in the placenta (Supplemental Table S4).

### Large EVs Are Enriched for Mitochondrial Transcripts

We then identified mitochondrial transcripts validated by MitoCarta 3.0, a curated literature-based human mitochondrial database and found that ∼10% (116/1134) of differentially abundant mRNAs in lEVs compared to sEVs were encoded by nuclear-encoded mitochondrial genes (Supplemental Table S4) [58]. This included transcripts encoding lactate dehydrogenase B (*LDHB*), translocase of outer mitochondrial membrane 22 (*TOMM22*), mitochondrial ribosomal protein subunits, PTEN induced kinase 1 (*PINK1*), voltage-dependent anion channel 1 and 2 (*VDAC1/VDAC2*), succinate dehydrogenase complexes (*SDH*), NADH:ubiquinone oxidoreductase subunits (*NDUF*), malate dehydrogenase 2 (*MDH2*), growth hormone inducible transmembrane protein (*GHITM*), and mitochondrial carrier 1 (*MTCH1*). The absence of mitochondrial-encoded genes in our sequencing data confirms that our Proteinase K treatment successfully eliminated intact mitochondria or non-encapsulated cellular debris co-isolated during EV preparation.

### Unique Transcripts Found Only in Large EVs

A subset of transcripts were detected in lEVs and not detected in sEVs (Supplemental Table S4). Notably, many stress-response transcripts such as aryl hydrocarbon receptor nuclear translocator (*ARNT)*, RNA binding motif protein 3 (*RBM3)*, activating transcription factor 4 (*ATF4)*, lactate dehydrogenase A (*LDHA)*, and selenoprotein P (*SELENOP)*, each induced by hypoxic, oxidative, and proteotoxic stress conditions [59–63]. An additional 18 nuclear-encoded mitochondrial genes annotated in MitoCarta 3.0 were detected only in lEVs (Supplemental Table S4). We note that transcript detection was substantially higher in lEVs overall, and lEV-exclusive detection may therefore reflect differences in detection sensitivity between sEVs and lEVs rather than selective packaging alone.

### Pathway Enrichment Analysis Highlights Pathways Important to Pregnancy

Ingenuity Pathway Analysis (IPA) of all differentially abundant transcripts revealed significantly enriched canonical pathways (padj < 0.05), providing a functional overview of EV-associated transcripts in early pregnancy (Figure 3e). Because 1,134 of the 1,142 differentially abundant transcripts were enriched in lEVs, the resulting pathway enrichment predominantly reflects lEV-associated transcripts. The eight sEV-enriched transcripts were insufficient for independent pathway analysis.

Cellular response to hypoxia was a top activated pathway, represented by 25 genes, consistent with the oxygen transition occurring across the gestational window sampled (11-13 weeks). Mitochondrial pathways were also identified, including oxidative phosphorylation (14 genes), and respiratory electron transport (12 genes). ESCRT pathway machinery was enriched in our dataset (7 genes), all of which were detected in lEVs and not sEVs. Additionally, hormone and pregnancy signaling pathways were enriched across differentially abundant transcripts including estrogen receptor signaling, which was among the top enriched pathways (46 genes), FLT3 signalin*g* (11 genes), and VEGF signaling (16 genes).

### Small RNA Levels Are Consistent in Small and Large EVs

EV-miRNAs has been extensively investigated in normal and complicated pregnancies [15,64–67]. However, a direct comparison of plasma-derived sEV and lEV small RNA composition from normal pregnancies remains limited. Here, we profiled the small RNA composition between these two EV-subtypes (Supplemental Figure S2). After rRNA removal, miRNAs and tRNA Derived RNAs (tDRs) each accounted for ∼25% of reads in both sEVs and lEVs (Figure 4a). PCA of miRNA profiles did not separate sEVs and lEVs (Figure 4b). Two sEVs samples, each from a different donor (ID 40 and 69), were distinct among the rest of the data points, however both had read depth and small RNA biotype distribution comparable to other samples (Supplemental Figure S2). With PC1 and PC2 explaining only 15% and 12% of the variance, this plot has limited power to resolve subtype differences.

**Figure 4.**
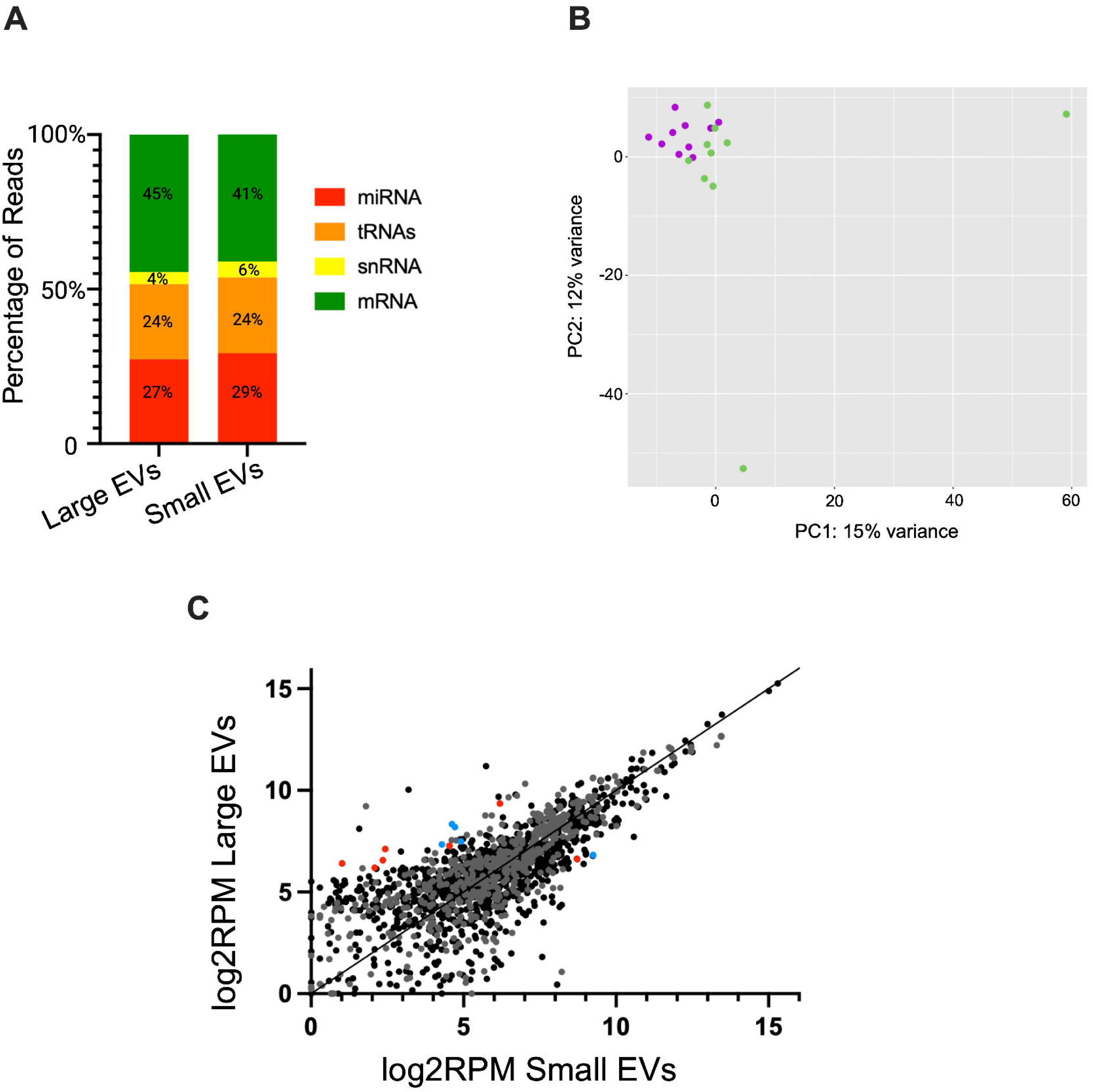
MiRNAs are consistently abundant in early pregnancy small and large EVs. (A) Average composition of total genome small RNA reads excluding rRNAs. (B) PCA of miRNA reads only of individual sEV and lEV samples. Purple dots represent lEVs and green represent sEVs. (C) Scatter plot of miRNAs and tRNAs normalized to total genome reads, highlighting differentially abundant miRNAs determined by DESeq2. Red dots represent miRNAs. Blue dots represent tRNAs. Significance threshold: log_2_FC ≥ |2| and adjusted *p*-value of ≤ 0.05. Created in BioRender.

Differential abundance testing identified 7 out of 1048 miRNAs that differed between sEVs and lEVs (Supplemental Table S5). lEVs were significantly enriched with 6 of the 7 differentially abundant miRNAs including hsa-mir-16-2, hsa-mir-223, hsa-mir-361, hsa-mir-527, hsa-mir-606, hsa-mir-942. hsa-mir-527 is a member of the primate-specific chromosome 19 microRNA cluster (C19MC) highly expressed by human placental trophoblasts and actively packaged into placenta-derived EVs [55,68–70]. Other C19 miRNAs detected in both sEVs and lEVs but not differentially abundant include hsa-mir-517a, hsa-mir-517b, hsa-mir-518b, hsa-mir-519a-1, and hsa-mir-519a-2, all of which have been exclusively expressed in the placenta (Supplemental Table S5) [69]. Additionally, lEVs were enriched with 4 tRNA-derived RNAs (tDRs): chr12.trna6-TrpCCA, chr2.trna10-SerTGA, chr2.trna21-SerTGA, and chr6.trna84-GlnTTG. In contrast, sEVs were enriched with hsa-mir-639 and chr1.trna64-GluTTC (Figure 4c).

Predicted mRNA targets of the differentially enriched miRNAs were generated using TargetScan and subsequently analyzed using IPA to identify enriched pathways associated with these target sets (Supplemental Table S5). We first used IPA to identify unique pathways enriched for the lEV-miRNA targets. Enrichment analysis highlighted pathways including cellular responses to hypoxia, electron transport chain complex assembly, MTOR signaling, and gluconeogenesis. Comparatively, targets associated with the sEV miR-639 related to tRNA and rRNA processing pathways.

### Canonical EV Protein Markers Associated with Small and Large EVs

To gain a comprehensive understanding of the sEV and lEV proteomic profile, EVs were isolated from nine out of the ten early pregnancy plasma samples for proteomic analysis using mass spectrometry. To enhance the robustness of our data, only proteins that were quantified in at least six out of the nine samples in either group were considered for analysis. After this correction, 1266 total proteins were identified. Notably, we found 29 proteins unique to lEVs and 58 unique to sEVs (Figure 5a)(Supplemental Table S6).

**Figure 5.**
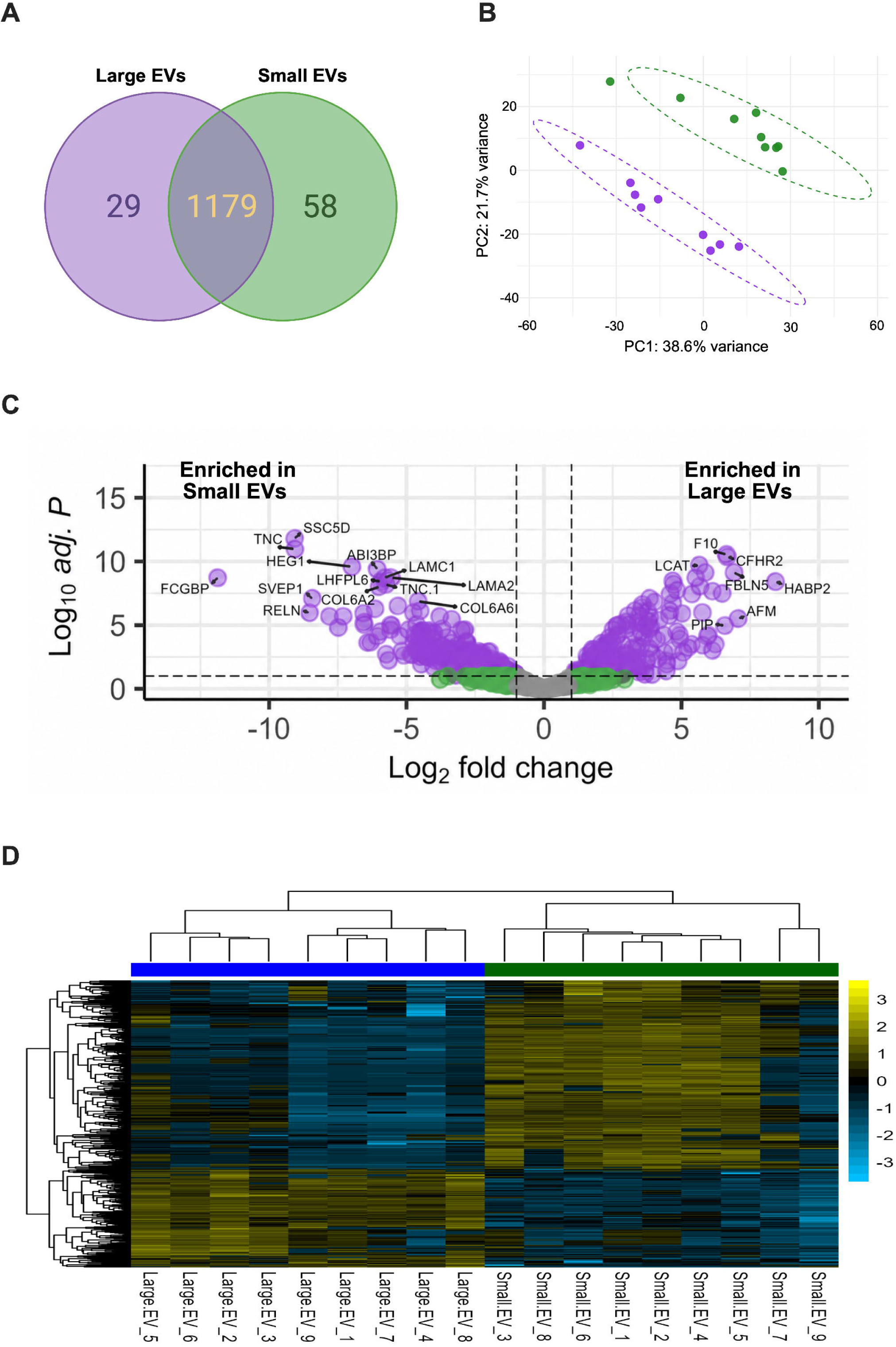
Small and large EVs have distinct proteomic profiles during pregnancy. (A) Venn diagram comparing protein hits between sEV and lEV samples.(B) Principal component analysis (PCA) of the proteome of individual sEV and lEV samples isolated from maternal plasma (11-15 weeks’ gestation). In total, there were nine lEV samples and nine sEV samples used for downstream analysis. (C) Volcano plot depicting log_2_ fold change and log_10_ adjusted *p*-value of identified proteins in sEVs and lEVs. Purple dots indicate differentially abundant proteins. A positive fold-change represents enrichment in lEVs, while a negative fold-change represents enrichment in sEVs. Significance threshold: log_2_FC ≥ |1| and adjusted *P*-value of ≤ 0.1. (D) Heatmap depicting the expression (log_2_ transformed) of differentially abundant proteins across EV subtypes. Created in BioRender.

Using the Vesiclepedia platform, we identified EV-specific proteins in our dataset to validate the accuracy of our EV enrichment preparations [71]. Out of the top 100 Vesiclepedia EV proteins, 65 overlapped with our sEV and lEV samples, including but not limited to CD9, CD81, and CD63 (Supplemental Table S6). We found that sEVs were enriched with top canonical EV markers, such as CD81, CD63, and ALIX, while lEVs were enriched with other EV proteins, including fibronectin 1 (FN1), gelsolin (GSN), and myosin-9 (MYH9) (Supplemental Table S7). Aligning with our immunoblot results, CD9 was also highly abundant in both EV subtypes via mass spectrometry.

### Enrichment of Placental and Mitochondrial-Related Proteins in Small and Large EVs

PCA revealed the distinct clustering of sEVs and lEVs, suggesting a unique proteomic profile between the two EV populations as previously reported (Figure 5b) [72–74]. To better understand the components that are driving EV subtype clustering during early pregnancy, we investigated the differentially abundant proteins between the groups. 549 proteins were considered differentially abundant, with 188 more abundant in lEVs and 361 more abundant in sEVs (Figure 5c)(Supplemental Table S6). Hierarchical clustering further illustrated the separation between EV subtypes. The heatmap visualized relative protein abundance across all samples grouped by sEVs and lEVs, revealing subtype-specific expression patterns (Figure 5d).

Similar to our transcriptomic analyses, we sought to identify differentially abundant proteins found in human placental proteomic data to reveal subtype-specific placental signatures (Supplemental Table S8) [75]. sEVs were differentially enriched for adiponectin (ADPIOQ), syndecan-1 (SDC1), nidogen-1 (NID1), pregnancy-associated plasma protein A (PAPPA), and endoglin (ENG). These proteins are associated with trophoblast basement-membrane structure, endocrine function, early placental vascular signaling, and nutrient transport [76–79]. In contrast, lEVs were differentially enriched for multiple PSGs, heparan sulfate proteoglycan 2 (HSPG2), collagen type XVIII alpha 1 chain (COL18A1), and CSH1, representing extracellular matrix components, syncytiotrophoblast-derived hormones, and maternal-fetal interface proteins [80–84].

Using MitoCarta 3.0, we then identified a panel of mitochondrial proteins, most of which were enriched in lEVs [58]. In total, 42 mitochondrial proteins were present in our dataset. Analyzing non-differentially abundant proteins, we identified L-2-hydroxyglutarate dehydrogenase (L2HGDH), catalase (CAT), peroxiredoxin 6 (PRDX6), superoxide dismutase 1 (SOD1) in both sEVs and lEVs (Supplemental Table S6). Comparing differentially abundant proteins, solute carrier family 44 member 1 (SLC44A1) was the only differentially abundant mitochondrial protein enriched in sEVs (Supplemental Table S9). Examples of proteins that were in more abundant in lEVs included chromosome 18 open reading frame 63 (C18orf63) and solute carrier family 25 member 5 (SLC25A5). In contrast, there were additional mitochondria proteins that were enriched in lEVs but in lower abundance including ATP synthase F1 subunit alpha (ATP5F1A), SOD2, thymidine kinase 2 (TK2), isocitrate dehydrogenase (NADP(+)) 2 (IDH2), and SLC25A3. These findings demonstrate the preferential enrichment of mitochondrial proteins in lEVs compared to sEVs despite sEVs having more proteins overall, further emphasizing the importance of size-fractionated, multi-omic profiling of plasma EVs.

### Multi-omic Integration Reveals Concordance Across mRNAs and Proteins

To evaluate concordance across molecular layers, we compared transcriptomic and proteomic datasets derived from our matched sEV and lEV samples. We first identified genes detected at both the transcript and protein level, yielding a shared subset of 367 RNAs and proteins (Supplemental Table S10). RNAs and proteins were classified as concordant when transcript and protein log2 fold changes followed the same direction, whereas discordance presented opposing directional changes. Directional agreement across mRNAs and proteins is visualized in a four-quadrant scatter plot (log2FC ≥ |1|) (Figure 6). Using these criteria, 47 genes demonstrated lEV concordance at the transcript and protein level including *VDAC2*, *SOD2*, *SLC25A3/5*, *PRDX6*, *MDH2*, and *PSG1*. In contrast, only 10 genes exhibited transcript and protein overlap in sEVs such as thioredoxin interacting protein (*TXNIP)*, radixin (*RDX)*, and sphingosine-1-phosphate receptor 1 (*S1PR1)*, genes associated with antioxidant response, cytoskeletal organization, and vascular signaling, respectively [85–87]. Collectively, these observations indicate a greater number of concordant features associated with lEVs compared to sEVs.

**Figure 6.**
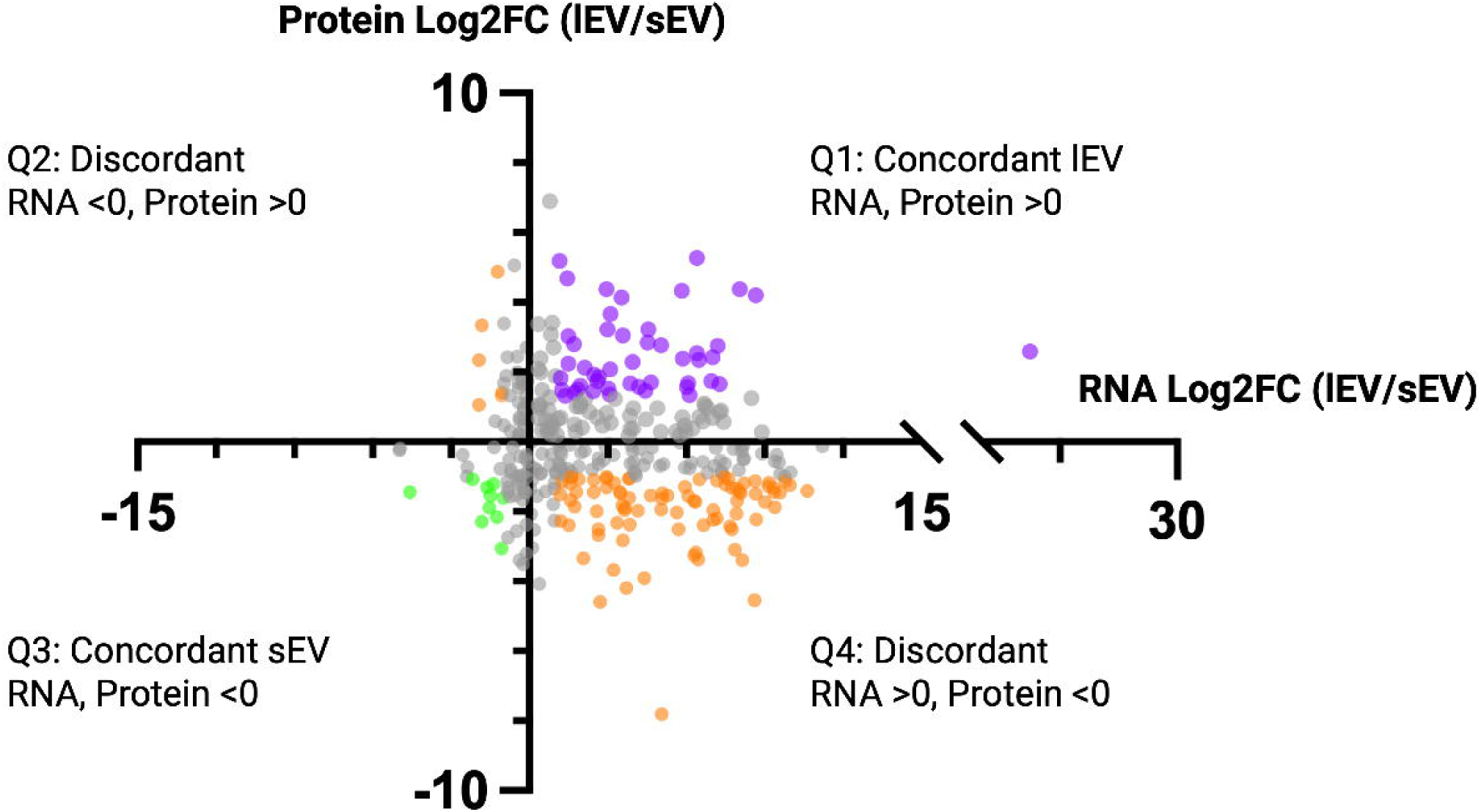
Multi-omic integration of small and large EVs reveals concordance across transcripts and proteins. A four-quadrant scatter plot of genes identified at both the transcript and protein level. Each dot represents individual genes plotted based on the average mRNA log_2_FC of lEV/sEVs (x-axis) versus the average protein log_2_FC of lEV/sEVs (y-axis). A positive fold-change represents enrichment in lEVs, while a negative fold-change represents enrichment in sEVs. Quadrant 1 indicates transcripts and proteins enriched in lEVs, with purple dots signifying a log_2_FC ≥ |1|. Quadrant 2 indicates transcripts enriched in sEVs and proteins enriched in lEVs. Quadrant 3 indicates transcripts and proteins enriched in sEVs, with green dots signifying a log_2_FC ≥ |1|. Quadrant 4 indicates transcripts enriched in lEVs and proteins enriched in sEVs. Orange dots represent discordant features with a log_2_FC ≥ |1| and grey dots represent features with a log_2_FC < |1|. Created in BioRender.

The remaining 207 genes showed opposite patterns in their transcript versus protein levels. For example, a gene might have high transcript levels in lEVs but high protein levels in sEVs, or vice versa. Among these 207 genes with opposite patterns, 92 showed large differences between lEVs and sEVs (log2FC ≥ |1|). Most commonly, the transcript was more abundant in lEVs while the protein was more abundant in sEVs, consistent with our overall finding that lEVs are associated with more transcripts while sEVs are associated with more proteins.

## Discussion

In this study, using size-fractionated plasma EVs and multi-omic profiling, we demonstrate that plasma-derived sEVs and lEVs carry distinct, but complementary molecular signatures during early pregnancy. Our data show that lEVs are predominately enriched for mitochondrial cargo, oxygen-sensing pathway modulators, endocrine and vascular signaling molecules, while sEVs are largely associated with extracellular matrix and basement membrane proteins. These differences were observed across different molecular layers and suggest that during early pregnancy lEVs provide more insight into systemic metabolism and cellular activity compared to sEVs. Together these data identify what circulating EV subtypes carry during uncomplicated pregnancy between 11-13 weeks, a window in which the placenta and maternal tissues undergo substantial physiological transition.

The most pronounced distinction between EV subtypes was the enrichment of mitochondrial cargo in lEVs across all multi-omic layers: mtDNA copy number, mitochondrial transcripts, and mitochondrial proteins spanning electron transport chain complexes, TCA cycle enzymes, and quality control machinery. This cargo is unlikely to represent co-isolated intact mitochondria. mtDNA was near-absent in EV-depleted samples, indicating that mtDNA detected is vesicle associated. Mitochondrially encoded transcripts were not detected and mitochondrial proteins were present in only low abundance whereas intact organelles would contribute both in large proportions given that the mitochondrial genome is actively transcribed. This pattern is more consistent with selective packaging of mitochondrial components into large EVs than co-isolated mitochondria [9–12,88].

The gestational window sampled in this study spans a critical placental shift from glycolysis to oxidative phosphorylation that supports trophoblast function, hormone production, and vascular remodeling [89]. Maternal tissues undergo coordinated adaptations over the same period, including adipocyte energy storage, immunometabolism to establish fetal tolerance, and vascular remodeling [90–94]. Enrichment of mitochondrial cargo in lEVs during this period is consistent with lEVs carrying material from metabolically active tissues, though our design cannot attribute it to a specific cellular source. Mitochondrial EV cargo may also have functional roles serving as both signaling molecules and metabolically active cargo [9]. In pregnancy, extravillous trophoblasts under oxidative stress release EVs with higher mtDNA content compared to normal pregnancies [47]. Outside of pregnancy, transfer of undamaged EV-mtDNA from healthy cells has been reported to rescue metabolically impaired recipient cells [13]. Mitochondrial proteins and intact mitochondria delivered by EVs have also been shown to incorporate into the mitochondrial network of recipient cells [9–12]. Whether the lEV mitochondrial cargo described here has comparable functional activity remains to be determined.

Another pronounced distinction between EV subtypes was the divergence in enrichment direction across proteins, miRNAs, and transcripts. Proteins were slightly more abundant in sEVs compared to lEVs (362 vs 188) and miRNAs were largely shared with only 7 differing between subtypes. Transcripts, however, were far more abundant in lEVs compared to sEVs (1,1134 vs 8). Three ‘omic analyses from the same donors using the same EV isolation techniques therefore produced three different patterns. Cargo sorting that favored lEVs would still be expected to yield a set of transcripts associated with sEVs, comparable to what was observed at the protein and miRNA level. This large discrepancy in the number of mRNAs enriched in lEVs compared to sEVs, despite small RNA libraries being prepared from the same total RNA pellet, suggests detection limits specific to sEV mRNAs that should be considered when interpreting such data. The enrichment of ESCRT transcripts among lEV cargo illustrates this point. ESCRT machinery is canonically associated with small EV biogenesis [18,20,23]. However, ESCRT proteins act within the parent cell to mediate vesicle formation, whereas ESCRT transcripts reflect RNA packaging rather than confirming ESCRT-dependent biogenesis. This discrepancy in addition to the frequency of transcripts and protein level discordance supports EV cargo sorting being governed by multiple selective pathways [2,18,30,95].

Within lEV cargo, the transcripts detected reflect the physiology of this gestational window. Detection of *ARNT*, encoding the HIF-1 beta subunit, alongside *CITED2*, a negative regulator, is more consistent with active modulation of the hypoxia response than with sustained hypoxia [89]. Estrogen receptor and VEGF signaling transcripts are consistent with the endocrine role of the placenta and with estrogen-driven upregulation of VEGF, while FLT3 signaling has been implicated in decidual immune function [96–102]. The enrichment of these pathways identify which transcript pathways are represented in lEV cargo during normal early pregnancy compared to sEVs. However, they do not establish active transcription in maternal or placental tissue.

EV-miRNAs are some of the most widely characterized EV cargo. In the context of pregnancy, many studies have associated various miRNAs with adverse pregnancy outcomes such as preeclampsia, preterm birth, gestational diabetes, and fetal growth restriction, positioning EVs as potential biomarkers [15,103–105]. In contrast with previous reports comparing EV subtypes, we did not find large differences in miRNA abundance between early pregnancy sEVs and lEVs despite pronounced transcript and protein differences [106,107]. Although two sEV samples appeared to be outliers by PCA, quality control assessment of read depth and alignment did not characterize those samples as statistical outliers. Those two samples may reflect the biological variation between each donor especially considering that the corresponding lEVs did not separate in the same pattern. We hypothesize that in this stage of pregnancy, miRNA-mediated signaling represents a shared functional role across EV populations, while major cargo differences primarily occur at the protein and transcript level, underscoring the value of multi-omic profiling.

The sEV fraction carried a distinct protein signature dominated by structural and extracellular matrix (ECM) proteins. ECM proteins are essential for the function of many cell types. During pregnancy, trophoblast cells actively remodel basement membranes during invasion, with extravillous trophoblasts degrading ECM to penetrate the decidua and syncytiotrophoblast cells organizing basement membrane components at the villous surface [108–111]. Simultaneously, maternal decidual stromal and endothelial cells secrete ECM and basement membrane proteins to support the implantation site and angiogenesis while maternal macrophages degrade ECM proteins during spiral artery remodeling [108,112,113]. Enrichment of these proteins in sEVs is consistent with remodeling activity across placental and maternal compartments, though cell-of-origin resolution would be required to attribute it. Notably, transcripts and their corresponding proteins were largely not co-enriched within the same subtype, indicating that the vesicle carrying a transcript is generally not the vesicle carrying its protein product, a pattern consistent with partly independent sorting of RNA and protein cargo across subtypes.

The relationship between EV size and cargo observed here sits within a broader spectrum of placental EV release. The human placenta sheds extracellular material across a wide size range, from exosomes and microvesicles to syncytial nuclear aggregates of 20 to 500 µm [53,54]. Differential centrifugation captures only part of this range, so our data describe two defined size fractions rather than the full spectrum of placental EV release. We detected transcripts and proteins overlapping published placental datasets, including cytotrophoblast and syncytiotrophoblast markers, consistent with a placental contribution to the circulating EV pool. We also detected several miRNAs specific to C19MC, including one which was differentially abundant between sEVs and lEVs, hsa-mir-527. C19MC is the largest human miRNA cluster that are highly expressed in the placenta, including some that are exclusive to the placenta. The presence of these miRNAs that overlap human placenta miRNA datasets in addition to mRNAs further support placental contribution to the total circulating pool. We do acknowledge however, that resolving true placental origin requires cell-specific based approaches. Although definitive EV cellular origin remains a technological constraint, approaches such as nanoscale high resolution flow cytometry and immunocapture using magnetic beads are being optimized for the isolation of placental EVs from maternal circulation [114–116].

A key advantage of multi-omic profiling is the ability to assess concordance and discordance across transcripts and proteins. Our integrated gene level analysis revealed predominantly discordant patterns at the molecule level, with transcripts enriched in lEVs while their corresponding proteins were enriched in sEVs. This systemic discordance is an important finding as EV RNA and protein sorting mechanisms are not well understood. While the overlapping features from our multi-omic gene-level analysis revealed modest concordance, there was convergence on shared biologically relevant pathways such as mitochondrial processes, vascular signaling, and stress response. We hypothesize that the co-enrichment of both transcript and protein within the same EV subtype likely reflects a regulated, and biologically relevant pathway in the originating cell. These concordant and discordant genes serve as baseline levels in normal pregnancy EV biology and highlights the capacity of EV subtypes to capture relevant signals. Our integrated multi-omic framework capturing concordance and discordance emphasizes the benefit of profiling EV subtype cargo across molecular layers. The extent to which EVs containing various cargo work together to influence maternal physiology needs to be investigated in functional studies.

While our data are striking, it is important to address the limitations of our work for accurate interpretation of the results. Our sample size of 10, while within the range of comparable EV multi-omic studies, limits statistical power to adjust for gestational age, maternal race, and fetal sex [117,118]. However, our matched size-fractionated design and comprehensive profiling across the same individuals enhances the confidence in our findings. Second, EV enrichment was confirmed by the identification of canonical EV proteins and vesicular morphology but complete depletion of Apo B100 was not achieved despite extensive processing of platelet-poor plasma. Therefore, residual lipoprotein may contribute to some detected proteins. Circulating EV mRNAs are frequently fragmented and present at low copy number, though fragmented transcripts can retain biological roles beyond translation including reflecting the transcriptional state of originating cells and serving as regulatory molecules stabilizing, localizing, and mediating translational activity of mRNAs in target cells [2–5][119–121]. In our mRNA data, we observed ∼70% of reads mapped to CDS exons and ∼26% to 3’ UTR exons (data not shown). This pattern enriched with coding sequences is consistent with functional protein translation, however validation studies are required to distinguish full-length from fragmented transcripts using methods such as RT-PCR spanning entire transcript lengths or long-read sequencing. Functional assays testing whether sEVs and lEVs differentially influence maternal immune cell phenotypes, endothelial cell function, or trophoblast gene expression would clarify if EVs serve solely as biomarkers or also as functional mediators.

We also recognize that full multi-omic profiling of both EV subtypes is unlikely to be practical as a screening strategy due to cost and clinical accessibility. We therefore view this approach as a discovery framework to characterize circulating EV subtypes rather than a candidate assay. We recommend that future biomarker studies include multi-omic profiling in a small, well-characterized cohort to identify a limited set of candidates. Then followed by validation in larger cohorts with targeted, lower-cost assays applied to the relevant EV subtype. Pairing this with surface marker isolation techniques would further allow biomarker candidates to be investigated with cell-of-origin resolution. Whether analysis in a defined EV subtype offers an advantage over bulk or whole-plasma measurement remains untested. sEVs typically dominate by particle count, but particle number does not equate cargo abundance.

This study establishes a multi-omic characterization of size-fractionated plasma EVs during early pregnancy. Through parallel profiling of matched sEVs and lEVs, we demonstrate that lEVs capture systemic metabolism, oxygen-sensing pathway modulation, endocrine and vascular signaling, and trophoblast activity, while sEVs reflect structural remodeling and organization in basement membrane tissues. The convergent enrichment of mitochondrial DNA, transcripts, and proteins in lEVs and not sEVs, revealed only through multi-omic profiling, validates the necessity of size-fractionation establishes baseline EV signatures in normal pregnancies.

## Acknowledgements

We would like to acknowledge the effort of the Clinical Research Coordinators at the Pregnancy and Perinatal Research Center at the Hospital of the University of Pennsylvania in enrolling patients and collecting samples. We also thank Dr. Luca Musante at the University of Pennsylvania’s Extracellular Vesicle Core for guidance in EV related methods. Thank you to Nicholas Williams and the Beckman Center for Cryo-Electron Microscopy for preparing the samples and cryo-EM support. Thank you to the Children’s Hospital of Pennsylvania Proteomics Core for your expertise in generating reliable EV proteomic data. Lastly, thank you to my co-authors and mentor Dr. Nicole Noren Hooten for your continued support throughout this process.

## Data Availability Statement

The RNA sequencing data reported in this study are deposited in GEO: GSE320491(mRNAseq) and GSE322470(miRNAseq). The mass spectrometry proteomics data have been deposited to the ProteomeXchange Consortium via the PRIDE partner repository with the dataset identifier PXD074792. All other data are available in the main text.

## Author Contributions

K.A.: conceived and designed research, analyzed data, performed experiments, interpreted results of experiments, prepared figures, drafted manuscript, edited and revised manuscript. T.H.: performed experiments and drafted manuscript. A.S.: analyzed data, interpreted results of experiments, prepared figures. E.M.: performed experiments. H.F., L.S.: conceived and designed research, analyzed data, interpreted results of experiments, prepared figures, edited and revised manuscript. R.L.: drafted manuscript. S.P., N.S., C.C., R.S.: conceived and designed research, interpreted results of experiments, edited and revised manuscript, approved final version.

