## Supplemental Figure S1 for "Multi-omic profiling defines subtype-specific extracellular vesicle signatures in early human pregnancy"

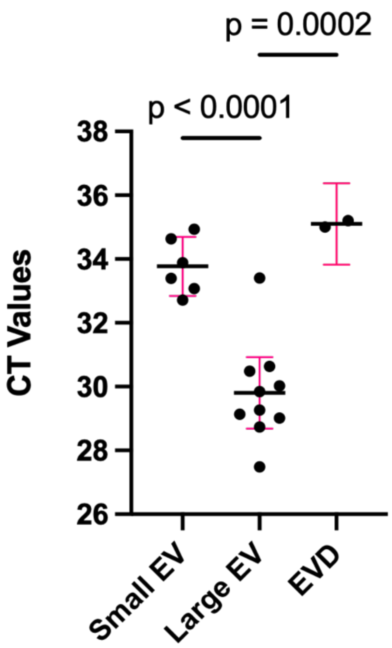


**Figure S1.** CT values per small and large EV sample and EV-depleted samples for mtDNA quantification.
