## Supplemental Figure S2 for "Multi-omic profiling defines subtype-specific extracellular vesicle signatures in early human pregnancy"

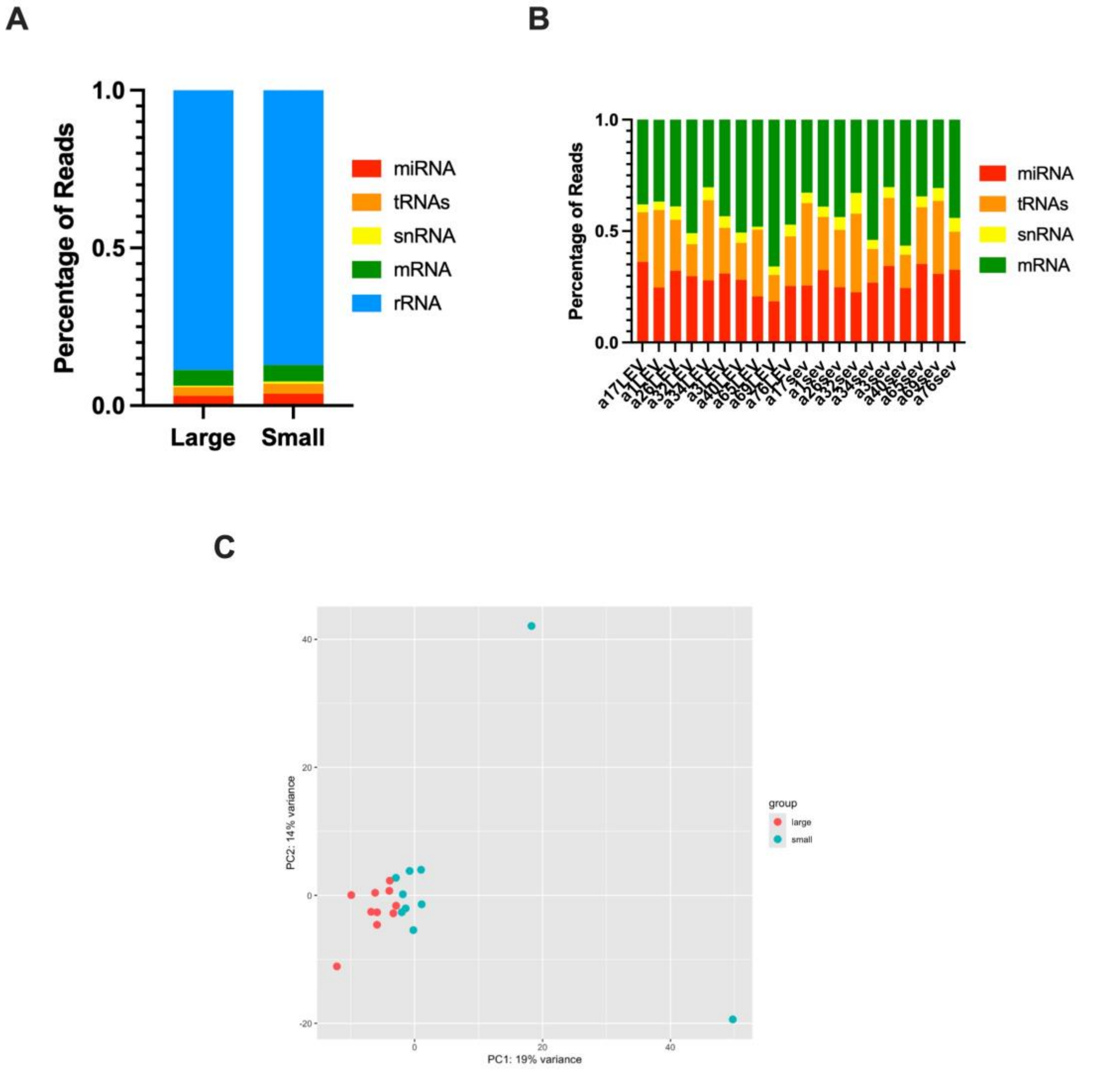


**Supplemental Figure S2.** (A) Small and large EV average composition of total genome small RNA reads. (B) Total genome reads across individual samples excluding rRNAs. (C) PCA of all small RNA species. Red dots represent large EVs and blue represent small EVs. Created in BioRender. https://BioRender.com/hmscv2x
