## Supplemental Table S1 for "Multi-omic profiling defines subtype-specific extracellular vesicle signatures in early human pregnancy"

| **Sample ID** | **Large EVs** | | **Small EVs** | |
|---|---|---|---|---|
| | Size (nm) | Concentration (particles/mL) | Size (nm) | Concentration (particles/mL) |
| **1** | 229.4 | 3.40E+09 | 89.5 | 4.00E+10 |
| **3** | 148.2 | 6.55E+09 | 91.3 | 3.60E+10 |
| **17** | 139.1 | 1.65E+10 | 87.9 | 3.15E+10 |
| **26** | 221.3 | 2.55E+09 | 107.2 | 2.25E+10 |
| **32** | 286.3 | 2.10E+09 | 130.7 | 5.87E+09 |
| **34** | 285.7 | 3.15E+09 | 123.1 | 5.40E+09 |
| **40** | 201.9 | 1.40E+09 | 128.3 | 4.65E+09 |
| **65** | 159.3 | 5.00E+09 | 80.4 | 1.45E+11 |
| **69** | 249.9 | 2.05E+09 | 114.4 | 3.53E+09 |
| **76** | 242.0 | 4.10E+09 | 138.2 | 2.40E+09 |
| **Average** | **216.3** | **4.68E+09** | **109.1** | **2.97E+10** |

**Supplemental Table S1.** EV size and concentration from 800 uL of platelet poor plasma from each subject. Samples used for mtDNA quantification.
