## Supplemental Table S2 for "Multi-omic profiling defines subtype-specific extracellular vesicle signatures in early human pregnancy"

| Sample ID | Large EVs | | Small EVs | |
|---|---|---|---|---|
| | Size (nm) | Concentration (particles/mL) | Size (nm) | Concentration (particles/mL) |
| **1** | 212.8 | 1.30E+10 | 91.3 | 3.35E+10 |
| **3** | 198.7 | 1.50E+10 | 87.9 | 4.30E+10 |
| **17** | 212.6 | 3.83E+10 | 103.4 | 3.35E+10 |
| **26** | 197.6 | 2.13E+09 | 89.6 | 1.55E+10 |
| **32** | 142.9 | 1.45E+10 | 89.0 | 8.30E+09 |
| **34** | 245.7 | 1.30E+09 | 123.8 | 1.20E+10 |
| **40** | 198.6 | 3.25E+09 | 141.4 | 3.25E+09 |
| **65** | 166.7 | 4.30E+10 | 83.3 | 2.00E+11 |
| **69** | 179.1 | 3.50E+09 | 96.4 | 2.10E+09 |
| **76** | 172.0 | 3.35E+11 | 87.9 | 1.15E+10 |
| **Average** | **192.7** | **4.69E+10** | **99.4** | **3.63E+10** |

**Supplemental Table S2.** EV size and concentration from 1 mL of platelet-poor plasma from each subject. Samples used for RNA sequencing.
