## Supplemental Table S3 for "Multi-omic profiling defines subtype-specific extracellular vesicle signatures in early human pregnancy"

| Sample ID | mtDNA Copies/LEV | mtDNA Copies/SEV |
|---|---|---|
| **1** | 1.54E-04 | 4.84E-05 |
| **3** | 1.35E-04 | 2.67E-05 |
| **17** | 1.01E-04 | - |
| **26** | 6.73E-05 | - |
| **32** | 1.45E-04 | 4.00E-05 |
| **34** | 7.27E-05 | 3.00E-05 |
| **40** | 9.21E-05 | - |
| **65** | 3.34E-04 | - |
| **69** | 1.65E-05 | 5.47E-05 |
| **76** | 1.78E-04 | 6.27E-05 |
| **Average** | **1.29E-04** | **4.38E-05** |

**Supplemental Table S3.** The number of mtDNA copies calculated per EV. 4E7 large EVs and 1E8 small EVs were used to isolate mtDNA. Samples with no measurable amount of mtDNA were reported as "-"and removed from analysis.
