## Supplemental Table S7 for "Multi-omic profiling defines subtype-specific extracellular vesicle signatures in early human pregnancy"

| ***Protein*** | ***Symbol*** | ***EV type*** | ***Average Intensity (log2)*** | ***q*** |
|---|---|---|---|---|
| *alpha-2-macroglobulin* | A2M | Small | 9.56 | 0.001 |
| *galectin-3-binding protein* | LGALS3BP | Small | 8.79 | <0.001 |
| *transferrin receptor* | TFRC | Small | 5.24 | 0.001 |
| *syndecan binding protein* | SDCBP | Small | 3.79 | <0.001 |
| *milk fat globule EGF and factor V/VIII domain containing* | MFGE8 | Small | 2.66 | 0.099 |
| *CD81 molecule* | CD81 | Small | 2.35 | <0.001 |
| *EH domain containing protein 1* | EHD1 | Small | 1.98 | 0.075 |
| *programmed cell death 6-interacting protein* | PDCD6IP (ALIX) | Small | 1.31 | 0.002 |
| *CD63 molecule* | CD63 | Small | 1.14 | 0.003 |
| *solute carrier family 3 member 2* | SLC3A2 | Small | 0.69 | 0.025 |
| *tumor susceptibility 101* | TSG101 | Small | -0.55 | 0.001 |
| *fibronectin 1* | FN1 | Large | 9.64 | 0.036 |
| *complement C3* | C3 | Large | 9.22 | <0.001 |
| *gelsolin* | GSN | Large | 5.16 | <0.001 |
| *myosin heavy chain 9* | MYH9 | Large | 3.02 | 0.073 |
| *actinin alpha 1* | ACTN1 | Large | 2.69 | 0.040 |
| *heat shock protein family A (Hsp70) member 5* | HSPA5 | Large | 1.80 | 0.047 |
| *actinin alpha 4* | ACTN4 | Large | -0.90 | 0.091 |

**Table S7.** Differentially abundant EV proteins. Average intensity represents the intensity across samples in a specific EV group. Negative $\log_2$ values reflect low-abundance proteins below the baseline intensity value. Adjusted *p*-value or $q<0.1$.
