## Supplemental Table S8 for "Multi-omic profiling defines subtype-specific extracellular vesicle signatures in early human pregnancy"

| *Protein* | *Symbol* | *EV type* | *Average Intensity (log2)* | *q* |
|---|---|---|---|---|
| *adiponectin, C1Q and collagen domain containing* | ADIPOQ | Small | 2.07 | 0.005 |
| *syndecan 1* | SDC1 | Small | 0.68 | 0.038 |
| *nidogen 1* | NID1 | Small | 0.63 | 0.057 |
| *pappalysin 1* | PAPPA | Small | -0.83 | <0.001 |
| *endoglin (CD105)* | ENG | Small | -1.55 | 0.091 |
| *fibronectin 1* | FN1 | Large | 9.64 | 0.036 |
| *pregnancy specific beta-1-glycoprotein 1* | PSG1 | Large | 4.05 | <0.001 |
| *pregnancy specific beta-1-glycoprotein 9* | PSG9 | Large | 3.12 | 0.021 |
| *pregnancy specific beta-1-glycoprotein 4* | PSG4 | Large | 2.18 | 0.007 |
| *heparan sulfate proteoglycan 2* | HSPG2 | Large | 0.75 | 0.007 |
| *collagen type XVIII alpha 1 chain* | COL18A1 | Large | 0.49 | 0.046 |
| *chorionic somatomammotropin hormone 1* | CSH1 | Large | -0.33 | <0.001 |
| *pregnancy specific beta-1-glycoprotein 6* | PSG6 | Large | -0.99 | 0.029 |

**Table S8.** Differentially abundant placental-related proteins crossed referenced with human placenta proteome. Proteins ordered by EV subtype and descending average intensity. Negative $log_2$ values reflect low-abundance proteins below the baseline intensity value. Adjusted *p*-value or q<0.1.
