## Supplemental Table S9 for "Multi-omic profiling defines subtype-specific extracellular vesicle signatures in early human pregnancy"

| ***Protein*** | ***Symbol*** | ***EV type*** | ***Average Intensity (log2)*** | ***q*** |
|---|---|---|---|---|
| *solute carrier family 44 member 1* | SLC44A1 | Small | 1.17 | 0.023 |
| *chromosome 18 open reading frame 63* | C18orf63 | Large | 3.04 | <0.001 |
| *solute carrier family 25 member 5* | SLC25A5 | Large | 0.42 | 0.020 |
| *ATP synthase F1 subunit alpha* | ATP5F1A | Large | -0.21 | 0.019 |
| *superoxide dismutase 2* | SOD2 | Large | -0.81 | 0.016 |
| *thymidine kinase 2* | TK2 | Large | -0.95 | 0.016 |
| *isocitrate dehydrogenase (NADP(+)) 2* | IDH2 | Large | -0.98 | 0.017 |
| *solute carrier family 25 member 3* | SLC25A3 | Large | -1.18 | 0.066 |
| *ATP synthase F1 subunit beta* | ATP5F1B | Large | -1.19 | 0.075 |
| *peroxiredoxin 3* | PRDX3 | Large | -1.26 | 0.015 |
| *microsomal glutathione S-transferase 3* | MGST3 | Large | -1.56 | 0.074 |
| *dynamin 1L* | DNM1L | Large | -1.58 | 0.081 |
| *voltage dependent anion channel 2* | VDAC2 | Large | -2.02 | 0.075 |
| *ATP synthase membrane subunit f* | ATP5MF | Large | -2.03 | 0.044 |
| *transmembrane protein 14C* | TMEM14C | Large | -2.50 | 0.015 |
| *mitochondrially encoded ATP synthase membrane subunit 6* | MT-ATP6 | Large | -2.60 | 0.077 |

**Table S9.** Differentially abundant mitochondrial-related proteins crossed referenced with the human placenta proteome and MitoCarta 3.0. Proteins ordered by EV subtype and descending average intensity. Negative $log_2$ values reflect low-abundance proteins below the baseline intensity value. Adjusted *p*-value or q<0.1.
